# Rodent Gut Bacteria Coexisting with an Insect Gut Virus in Parasitic Cysts: Metagenomic Evidence of Microbial Translocation and Co-adaptation in Spatially-Confined Niches

**DOI:** 10.1101/2024.03.22.585885

**Authors:** Amro Ammar, Vaidhvi Singh, Sanja Ilic, Fnu Samiksha, Antoinette Marsh, Alex Rodriguez-Palacios

## Abstract

In medicine, parasitic cysts or cysticerci (fluid-filled cysts, larval stage of tapeworms) are believed to be sterile (no bacteria), and therein, the treatment of cysticerci infestations of deep extra-intestinal tissues (*e*.*g*., brain) relies almost exclusively on the use of antiparasitic medications, and rarely antibiotics. To date, however, it is unclear why common post-treatment complications include abscessation. This study quantified the microbial composition of parasitic cyst contents in a higher-order rodent host, using multi-kingdom shotgun metagenomics, to improve our understanding of gut microbial translocation and adaptation strategies in wild environments. Analysis was conducted on DNA from two hepatic parasitic cysts (*Hydatigera (Taeenia) taeniaeformis*) in an adult vole mouse (*Microtus arvalis*), and from feces, liver, and peritoneal fluid of three other vole family members living in a vegetable garden in Ohio, USA. Bacterial metagenomics revealed the presence of gut commensal/opportunistic species, including *Parabacteroides distasonis, Klebsiella variicola, Enterococcus faecium*, and *Lactobacillus acidophilus*, inhabiting the cysts. *Parabacteroides distasonis* and other species were also present outside the cyst in the peritoneal fluid. Remarkably, viral metagenomics revealed various murine viral species, but unexpectedly, it detected an insect-origin virus from the army moth (*Pseudaletia*/*Mythimna unipuncta*) known as Mythimna unipuncta granulovirus A (MyunGV-A) in both cysts, and in one fecal and one peritoneal sample from two different voles, indicating survival of the insect virus and adaption in voles. Metagenomics also revealed a significantly lower probability of fungal detection in the cysts compared to other samples (peritoneal fluid, p<0.05; and feces p<0.05), with single taxon detection in each cyst for *Malassezia* and *Pseudophaeomoniella oleicola*. The samples with a higher probability of fungi were the peritoneal fluid. In conclusion, commensal/pathobiont bacterial species can inhabit parasitic tapeworm cysts, which needs to be considered during therapeutic decisions of cysticerci or other chronic disease scenarios where immune privileged and spatially restricted ecosystems with limited nutrients and minimal presence of immune cells could facilitate microbial adaptation, such as within gut wall cavitating micropathologies in Crohn’s disease.

## Introduction

The microbial communities, with their diversity and interactions, continue to intrigue scientists due to the unforeseen complexities of symbiosis in many ecosystems^1,2^, including spatially-confined niches such as within parasitic cysts. The interplay of microbial populations is critical in maintaining homeostasis and health in the mammalian gut ecology. Therefore, the study of these microbial communities could help us determine what influences these dynamic relationships in restricted ecological niches^3,4^, or the evasion of the host immunity in other confined biological niches, such as within gut wall-associated cavitating micropathologies in Crohn’s disease^5,6^, which we found to have genomic evidence for niche specific genomic exchange that could favor silent commensalism and immune evasion^6-9^.

The relevance of cysticerci in medicine is associated with the diseases that they may induce as they travel through the body and stop to enter into their seemingly quiescent cystic stage, where they become space-occupying masses that gradually develop unnoticed by affected individuals, and which could drag bacteria on integumentary micro-cavitations, as seen with electron microscopy on tapeworms^10^, from the gut or ingested foods within the gut as parasitic larvae migrate. Of immunological interest, such migratory parasites enter in contact with tissues, triggering host immunity which could presumably stress and drive the selection of microbial communities that successfully survive evading the immune system. How these communities are assembled in areas distant from the gut where there is a narrow range of nutrient sources and stressors, and where there is no physical removal of bacteria by peristalsis remains unknown.

Commonly, parasitic tapeworms in domestic animals in the adult forms live in carnivorous species, which then develop cysts within the peritoneal cavity in intermediate hosts, including humans, as larva migrate after ingestion and activation within the intestinal tract^11-13^. *Echinococcus granulosus* and other parasites alike^12^, including *Taenia* and *Hydatigera*, are one of the many pathogenic parasites that seem to facilitate intricate interactions within their intermediary hosts during migration at the larva stage^12^. In addition to helping us determine to what extent the cysts could contain microbial species that may contribute to our understanding of how to best treat, for instance, brain cysticerci in humans, especially children who are almost always affected by one cysticercus^14^, the study of parasitic cysts in rodents provides an attractive model^15^ for studying microbial survival and symbiosis away from the gut and its nutrient-rich dietary and fecal environment.

Herein, we report the results from a metagenomics community composition analysis in various tissue samples from a family of wild voles, one of which was affected with two extra-hepatic parasitic cysts that resembled the appearance of *Echinococcus granulosus* cysts common in 80.5% of affected humans^14^ (dimensions10 and 12 mm diameter), but was confirmed as *Hydatigera* (formerly *Taeniea*) in this study. The purpose of this study was to assess and report the community composition metagenomics analysis of parasitic systems in the context of other organs (feces, liver, peritoneal fluid) among family members of the vole affected with the cysts to catalogue the bacterial communities that may translocate within parasitic larval stages, and thrive outside of the typical gut milieu, in the parasitic cysts.

## Methods

### Animals and location

Common voles (*Microtus arvalis*) were located in an experimental community vegetable garden, in peri-urban Ohio, and trapped as a part of a pest control program using over-the-counter approved humane mouse traps. Samples were collected in the field, transferred to the lab, frozen, and processed for DNA extraction. The study included a total of eleven tissue samples obtained from four different voles. The first vole provided the two cysts located in its abdominal cavity, anchored on the surface of the liver capsule; no other samples were collected. For the other three animals, we sampled liver, peritoneal fluid, and feces. The samples were collected and frozen for metagenomic analysis which accounted for bacteria, viruses, and fungi.

### Identification of the parasitic cysts

The frozen cystic-like structure and the cyst fluid DNA samples were sent overnight to the Diagnostic Veterinary Parasitology Laboratory, at the Veterinary Medical Center, The Ohio State University. The cyst was thawed briefly, extraneous host tissue removed from the cream/white colored cyst using small forceps and a needle. The cyst was placed on a microscope, covered with a coverslip and pressure applied to flatten the structure for photo microscopy. Images were captured using an Olympus BX41 with CellSens software.

### Visualization and anaerobic culture of fluid from the parasitic cysts

To help determine whether the cysts were harboring live bacteria, we visualized the liquid in phase contrast medium using a 1000x magnification, and also cultured^16^ the fluid by spread-plating 20 microliters onto a pre-reduced tryptic soy 5% defibrinated sheep blood agar (80% N–10% H– 10% CO2, at 37°C; Thermo Fisher Scientific) using a variable-atmosphere anaerobic Whitley workstation A85 (540 plate capacity; Microbiology International, Inc.) as described^6,9^. Individual colonies were sub-cultured and purified in the same agar and then immediately identified using matrix-assisted laser desorption ionization–time of flight [MALDI-TOF] mass spectrometry) and banked in pre-reduced brain heart infusion broth with 7% dimethyl sulfoxide^6,9^.

### DNA extraction and metagenomics

DNA extraction was conducted using the DNAeasy Qiagen kit, while the DNA was quantified using the GloMax Plate Reader System (Promega) using the QuantiFluor® dsDNA System (Promega) chemistry. Samples were submitted for metagenomic analysis to a third party (CosmoID) which has validated methods and software^17,18^ for library preparation, sequencing and cloud-based computing for data analysis^19^.

### Library Prep and Sequencing

DNA libraries were prepared using the Nextera XT DNA Library Preparation Kit (Illumina) and IDT Unique Dual Indexes with total DNA input of 1ng. Genomic DNA was fragmented using a proportional amount of Illumina Nextera XT fragmentation enzyme. Unique dual indexes were added to each sample followed by 12 cycles of PCR to construct libraries. DNA libraries were purified using AMpure magnetic Beads (Beckman Coulter) and eluted in QIAGEN EB buffer. DNA libraries were quantified using Qubit 4 fluorometer and Qubit™ dsDNA HS Assay Kit. Libraries were then sequenced on an Illumina HiSeq X platform 2x150bp.

### Bioinformatics Analysis and metagenome classification

The system used utilizes a high-performance data-mining k-mer algorithm that rapidly disambiguates millions of short sequence reads into the discrete genomes engendering the particular sequences. The methodology employed in this study uses the CosmosID-HUB for fast and precise metagenomic analysis of microbiome data^18-20^, incorporating detection capabilities at the strain level across multiple kingdoms, along with antimicrobial resistance/virulence factors (AMR/VF), and functional analysis within a singular processing framework, which has been shown recently to perform well compared to other pipelines^17,18^. Herein, we solely report the microbial community composition since the interpretation of functional data for *Bacteroidota* has some limitations and complexity that we recently determined and which are under investigation^8^. The complete documentation for the analysis^19^ is available at https://docs.cosmosid.com/docs/methods (accessed March 20, 2024). This analysis platform is powered by three core components, a genBook database, a Kepler algorithm, and machine learning filters. The GenBook is a meticulously curated Multi-Kingdom Reference Database featuring over 180,000 genomes and gene sequences from bacteria, fungi, viruses, phages, and protists. Its curation process is designed to enhance sensitivity by reducing redundancy and ensuring homogeneity, particularly in densely populated clades such as *Staphylococcus aureus*. The database universal curation approach allows for consistent analysis across various sample types within a project, ensuring accuracy through genome quality control and minimizing false positives. The Kepler Algorithm is a patented, k-mer based algorithm that offers efficient and highly accurate profiling. It utilizes unique and shared kmers across the phylogenetic tree for precise near-neighbor placement, ensuring these kmers are phylogenetically stable and do not overlap with mobile genetic elements or the human genome. This approach, coupled with GenBook phylogenetic ontology, permits accurate differentiation down to the strain level. Lastly, the system uses Machine Learning Filters within the analysis pipeline which is enhanced by machine learning algorithms trained on over 10,000 samples, allowing for the distinction between genuine signals and background noise. This maintains high sensitivity and precision, as evidenced by superior F1 scores in benchmarks and community challenges.

For metagenomic analysis, whole genome shotgun sequencing data (in fastq or fasta formats) is used. Paired-end files may be combined for analysis if uploaded simultaneously. Following sample upload, the CosmosID-HUB automatically processes and generates detailed reports, including tables and visualizations for genome and gene databases, covering bacteria, fungi, protists, viruses, respiratory viruses, antimicrobial resistance, and virulence factors, which are shown in this report. As for performance evaluation, studies have validated CosmosID leading accuracy and resolution in detection. The system demonstrates exceptional identification accuracy across all taxonomic levels in benchmark datasets, significantly outperforming other tools, especially in sub-species and strain-level classification.

The metagenomics pipeline has two separable comparators, the first consists of a pre-computation phase for reference databases and the second is a per-sample computation^19^. The input to the pre-computation phase are databases of reference genomes, virulence markers and antimicrobial resistance markers that are continuously curated and added to an updated taxon database. The output of the pre-computational phase is a phylogeny tree of microbes, together with sets of variable length k-mer fingerprints (biomarkers) uniquely associated with distinct branches and leaves of the tree.

The second per-sample computational phase searches the hundreds of millions of short sequence reads, or alternatively contigs from draft *de novo* assemblies, against the fingerprint sets. This query enables the sensitive yet highly precise detection and taxonomic classification of microbial NGS reads. The resulting statistics are analyzed to return the fine-grain taxonomic and relative abundance estimates for the microbial NGS datasets. To exclude false positive identifications the results are filtered using a filtering threshold derived based on internal statistical scores that are determined by analyzing a large number of diverse metagenomes^19^.

### Metacestode DNA extraction and sequencing

Genomic DNA was extracted using DNeasy Blood & Tissue Kit (Qiagen) following manufacturer’s instructions for tissues with a slight modification. During the proteinase K digestion, the sample was continuously rotated at 56 C for 45 min. The mitochondrial 12S rRNA gene region PCR was targeted using 20 to 45 ng of genomic DNA per reaction along with Applied Biosystem Power SYBR Green PCR Master Mix and previously described primers, Cest F: 5’ AGTCTATGTGCTGCTTAT 3’ and Cest R: 5’ CCTTGTTACGACTTACCT 3. Oberli et al., 2023. Cycling consisted of 95C for 2 minutes followed by 50 cycles of 95C for 15 sec, 45C for 30 sec and 60C for 1 min on an Applied Biosystems StepOne Instrument. Control DNA of *Echinococcus granulosus, E. multilocularis* and *Taenia* sp. was provided by Kamilyah R. Miller (Kansas State University). The amplicon obtained from the cystic structure DNA from 6 different reactions were pooled and purified using QIAquick PCR Purification Kit spin columns. The purified product and primers were submitted to Genewiz for DNA sequencing. Two replicate experiments representing both forward and reverse were used to construct the consensus sequence. The 176 base pair DNA sequence was compared to published sequences using a nucleotide Blastn search (ncbi.nlm.nih.gov). The resulting DNA sequence was submitted to GenBank accession PP477764.

### Statistics

This report is primarily descriptive because the number of animals and samples tested were limited. Univariate analysis of metagenomic community composition and frequency statistics (presence/absence) for species of interest across the samples^21^ was conducted to determine if findings were random or significantly different from random. For this purpose, we used Fisher’s exact, or Chi-square statistics using GraphPad (v10.2.1) depending on the number of observations in each cell in a 2xn tables. Statistical significance for expected vs observed was held at p<0.05.

### Data availability

The metagenomic sequences and fastq files have been deposited in NCBI GenBank under the BioProject number PRJNA1053337. Entitled ‘gut microbiome that evades host immunity in wild rodents (vole) and parasitic cysts’, this project has 11 associated BioSamples and Sequence Read Archive (SRA) numbers for sharing with the scientific community under submission SUB14073752, and accessions SRR27223102 through SRR27223112, scheduled for release on April 18, 2024.

## Results

### Nucleotide sequence analysis of metcestode reveals *Hydatigera taeniaeformis*

An overview of the cysts and fecal, peritoneal fluid and liver samples, collected from 4 mice, and processed in this study can be found in **Figures 1A-D**. Although initial environmental assumptions suggested that *Echinococcus granulosus* was the most likely parasite (*e*.*g*., the presence of coyotes and other carnivores in the farm where the vegetable garden was implemented), DNA amplification and sanger sequencing results revealed the parasite metacestode was *Hydatigera taeniaeformis*, which forms a strobilocerus as its metacestode stage in the intermediate rodent host. The NCBI Blastn search showed the greatest percent identity (99%) to *Hydatigera* sp. (Genbank LC008533.1). Microscopic examination of the parasite confirmed that the segmentation patterns observed on the surface of the organism are suggestive of an immature strobilocercus metacestode stage which was discernable on the photomicrographs using the magnification and dorsal ventral flattening of the cyst. The lack of hooks and size suggests that this cyst-like structure is an immature strobilocercus.

**Figure 1.**
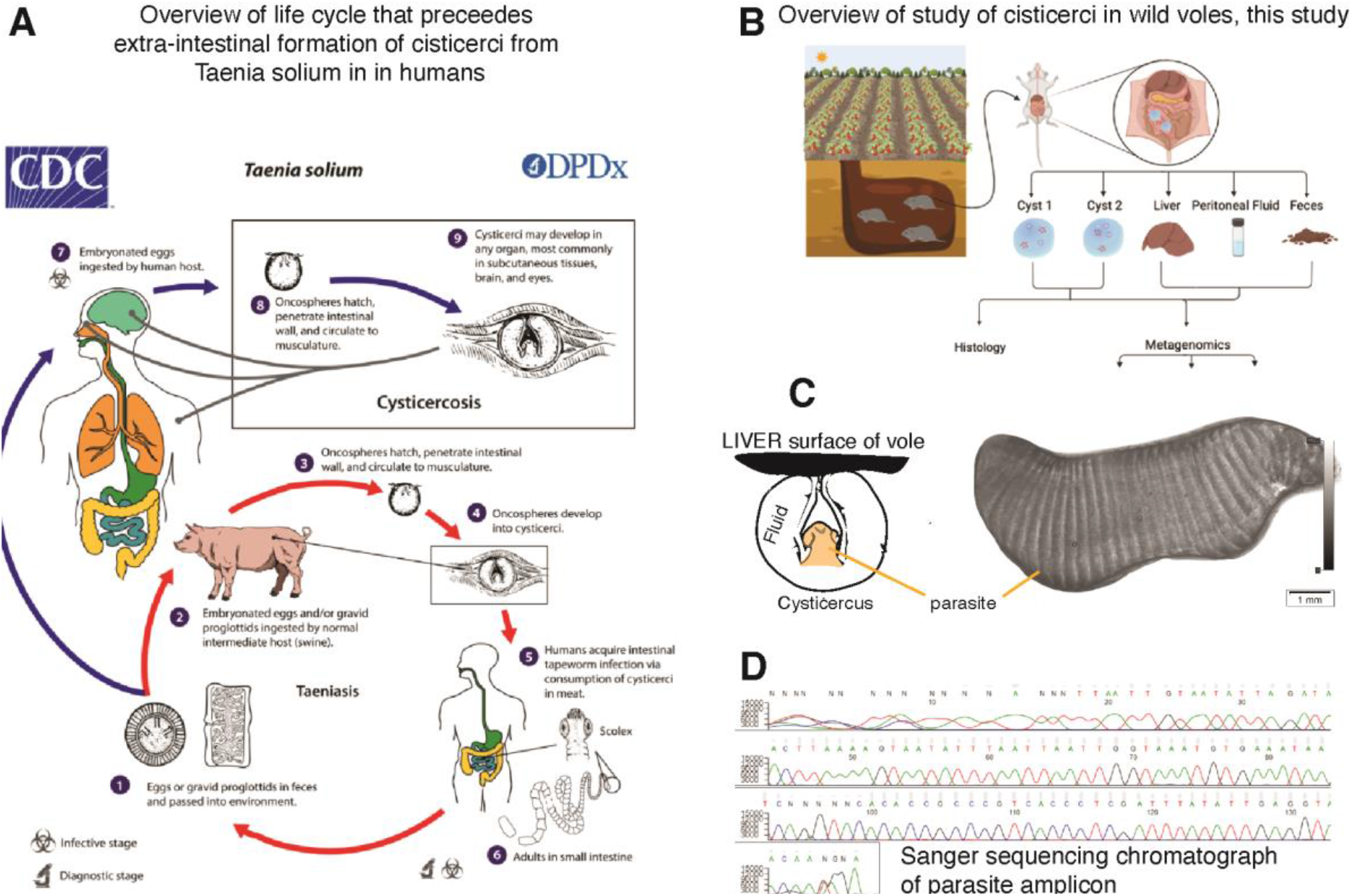
Overview of family of voles in this study and identification of the hepatic parasitic cyst as the larval stage of *Hydatigera taeniaeformis* tapeworm. **A)** Contextualization of the clinical and ecological relevance of human cysticercosis, exemplified with *Taenia solium*. (CDC public domain image). **B)** Samples collected from the voles in this report. **C**) Schematic of origin of samples within metacestode (parasitic-cystic structure) for analysis and photomicrograph, illustrating distinctive microscopic segmentation of the parasite as indicative of an immature strobilocerus metacestode. Not detailed the parasite lacks visualized hooks and the overall size suggests that this cyst-like structure is an immature strobilocerus. **D**) Chromatogram after Sanger sequencing used for tapeworm identification as *Hydatigera taeniaeformis* illustrates pure DNA in the cyst samples tested in this study.

### Visualization and cultivation of fluid yielded *Enterococcus faecium*

Of interest, we were able to visualize the presence of highly mobile bacterial-like structures in the fluid examined under contrast phase microscopy and visualized a complex array of gram-positive and gram-negative bacteria. However, also of interest, cultivation of the cysticercus fluid only revealed the presence of pure colonies of *Enterococcus faecium*, which were identified using Maldi-Toff. Although gram-staining of biological samples do not resemble the textbook gram-stain description of microbes isolated on agar surfaces, the isolation of pure *Enterococcus* on the agar (typically gram-positive cocci), and not of other bacteria, indicates that the cohabitation of multiple bacteria in a spatially-confined nutrient-depleted biological niches, such as the cysts, could be rendering *Enterococcus* species more symbiotic with other community members, instead of being inhibitory once it is growing on an artificial nutrient rich medium such as 5% sheep blood TSA plates as we previously documented for a fecal *Enterococcus* strain against a co-inhabitant *Lactobacillus* in the intestinal tract in a mouse model of Crohn’s disease^22^. Metagenomic analyses where therein pursued to better characterize the non-cultivable species in the cysts.

### Metagenomics of the cystic fluid revealed gut commensal/opportunistic bacteria

Metagemomic analysis revealed a relatively simple bacterial community inside the two cysts, demonstrating that these symbiotic bacteria could avoid the immune system and flourish over time in a nutrient-depleted lesion. At the species level, *Klebsiella variicola* comprised 18.34% and 35.48% of the total bacterial population in cysts 1 and 2, respectively, followed by *Enterococcus faecium* in cyst 2 (32.59%). *K. variicola* was also highly abundant in the peritoneal fluid samples of vole 2 (33.34%), vole 3 (39.86%), and vole 4 (74.9%), and feces samples of vole 2 (29.66%) and vole 4 (37.96%). In contrast, the vole 3 liver was entirely inhabited by *Propionibacteriaceae*, which only constituted 8.95% of cyst 1 (**Figures 2A-B**).

**Figure 2.**
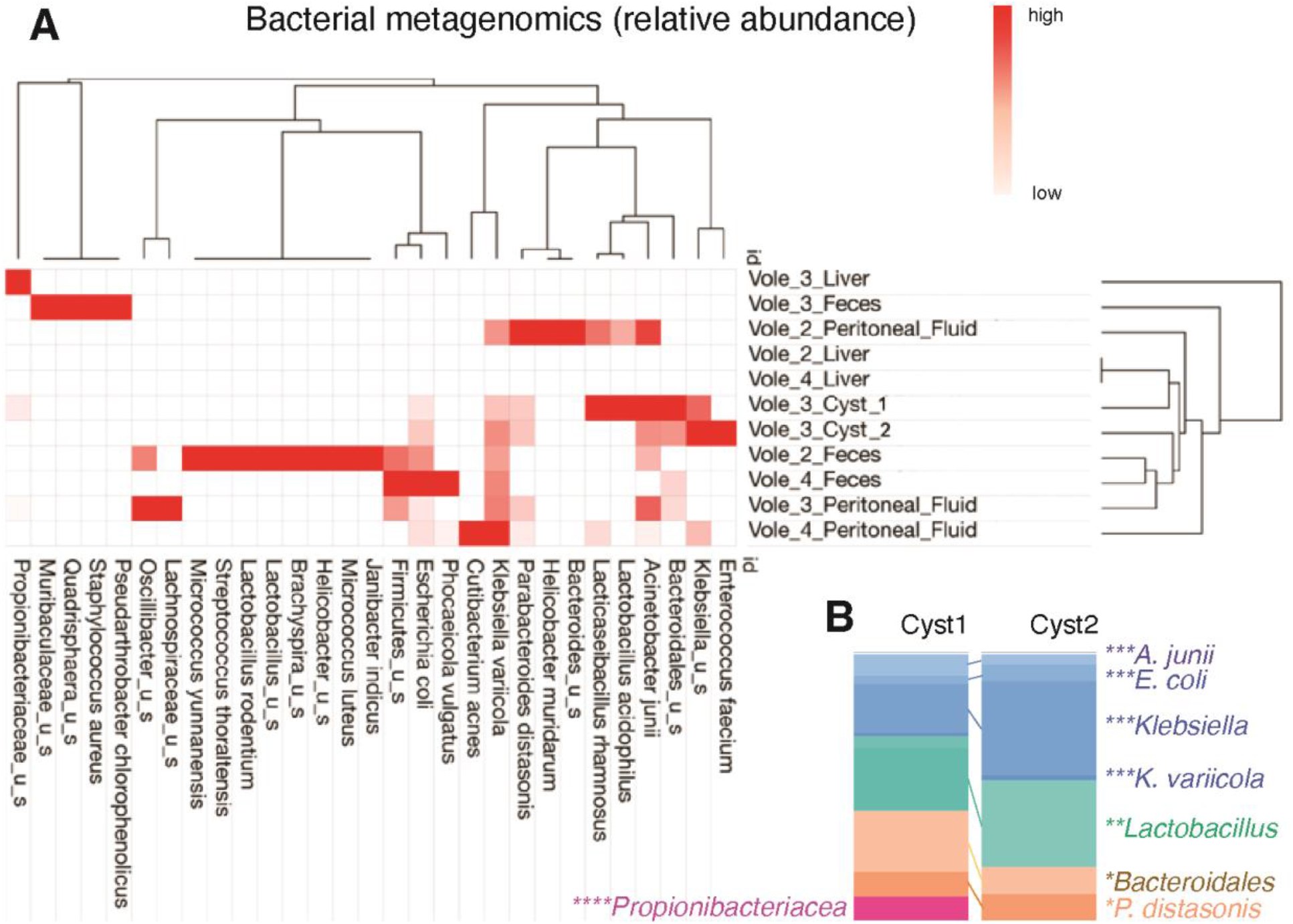
Metagenomic analysis identifies reproducible commensal *Bacteroidota* and opportunistic *Pseudomonadota* (*Enterobacteriaceae*) in the parasitic cysts. **A)** Relative abundance across samples. **B)** Comparison of bacteria in both cysts demonstrates consistent pattern of abundance among similar bacteria, including *K. variicola, P. distasonis*, and *Bacteroidales*.

*Bacteroidales* amassed an abundance of 23.29% in cyst 1 and 10.28% in cyst 2. At the species level, the cysts 1 and 2 had a very comparable abundance of *Parabacteroides distasonis* with 9.08% and 9.88%, respectively. The analysis of peritoneal fluid revealed *P. distasonis* as the most prevalent bacteria (42.39%) in vole 2, and as a highly abundant bacteria in vole 3 (9.36%). Of note, vole 3 was inhabited by *Quadrisphaera sp. DD2A* (44.69%). The latter finding is of notoriety, since the DNA of fecal samples in this study did not reveal a large number of bacterial taxonomic units, as expected from other studies we have conducted in human colonoscopy content and in mice^7,22^, with the same methodology, or using 16S rRNA microbiome studies^23,24^. This finding could be attributed to completely different gut microbiome in these wild animals who inhabit subterraneous environments and have different diets. Repeated testing confirmed the limited detection of OTUs in feces on this study.

### Identification of insect virus outside its natural habitat

Our metagenomic analysis reports, for the first time, the presence of a virus, Mythimna unipuncta granulovirus A (MyunGV-A), a virus adapted to the insect armyworm *Mythimna unipuncta*^25^ (which feeds on crops, including corn^26^), detected outside of its natural insect habitat, inside parasitic cysts within voles. MyunGV-A naturally infects and replicates within the larvae of the armyworm moth, *Mythimna unipuncta*, primarily within the cells of the midgut epithelium.

Our study identifies the presence of MyunGV-A in high abundance, suggesting that the virus may be thriving inside the cyst through cohabitation with the bacteria, and the absence of other viruses that were identified in the liver, peritoneum and feces of the other voles. MyunGV-A was found to be the only viral species in both cysts, while it was combined with Human mastadenovirus C in vole 2 peritoneal fluid (61.19%; **Figures 3A-B**). In vole 4 peritoneal fluid and vole 2 feces, MyunGV-A relative abundance was 100%, since no other viruses were detected. The liver samples were mostly inhabited by Moloney murine sarcoma virus and Murine osteosarcoma virus, which are expected infectious viruses of rodents. In the peritoneal fluid of vole 3, Abelson murine leukemia virus was the most prevalent (51.19%), followed by MyunGV-A (31.72%). This discovery indicates that MyunGV-A has probably adapted to voles, the parasite, or to the gut microbiome of mice, or that the microbiome provides metabolites that enable the virus to colonize other species (mouse or hydatygera), raising questions about the role of newly adapted, or transiently infecting baculoviruses in rodents and the implications for both the parasite and its mammalian host.

**Figure 3.**
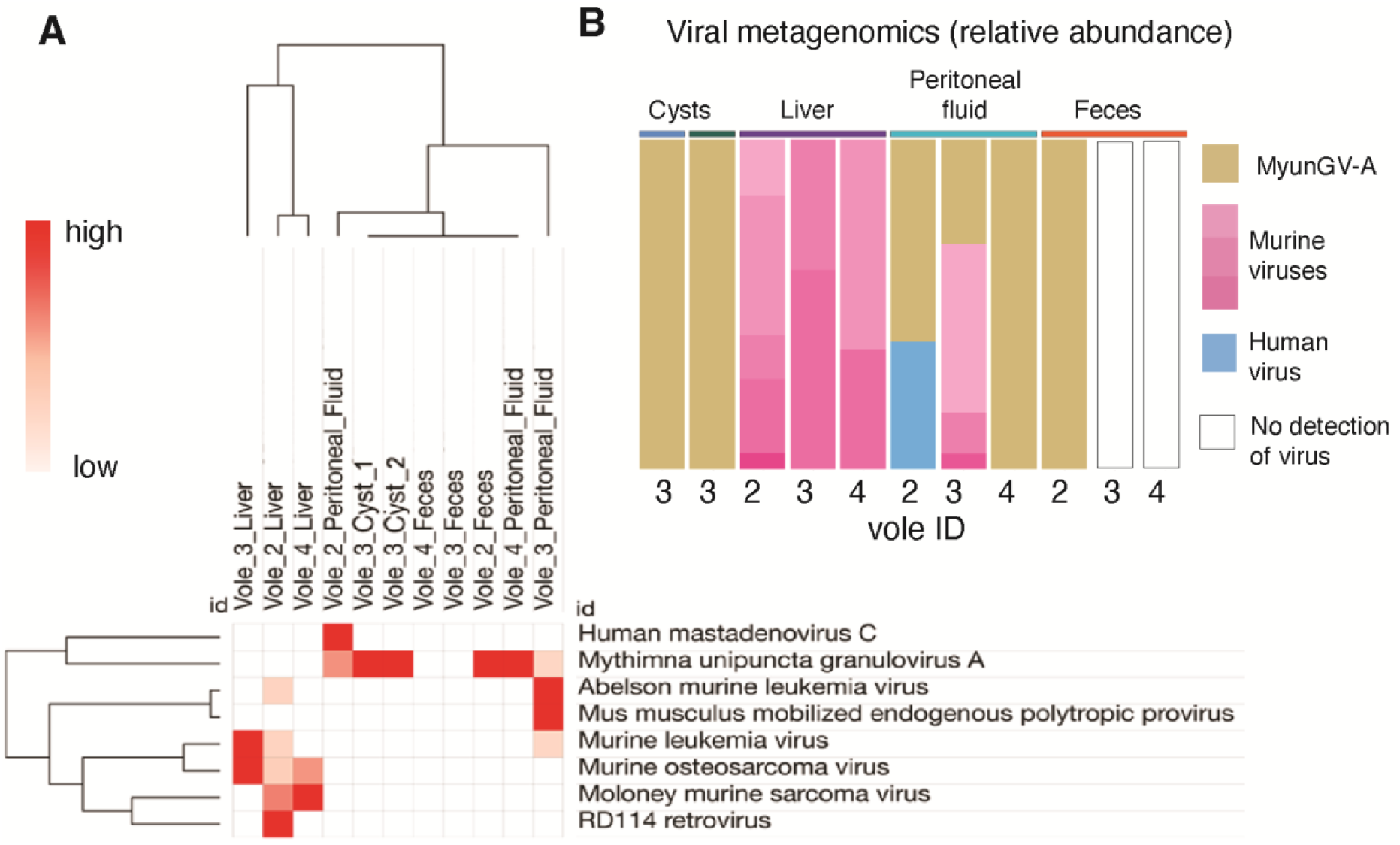
Metagenomic analysis of viruses revealed MyunGV-A in Hydatygera larval cysts and peritoneal fluid of wild voles. **A)** Hierarchical clustering of samples based on abundance. Euclidean distances. **B)** Abundance bar plot. MyunGV-A in both cysts was 100%, with similar findings in the peritoneal fluid and feces of two voles.

### Metagenomics suggests lower probability of fungal detection in the cysts

A binary analysis of the presence or absence of fungal DNA in the samples, regardless of the species identified, revealed that the animals had a significantly lower probability of fungal detection in the cysts compared to other samples (peritoneal fluid, p<0.05; and feces p<0.05), with single taxon detection in each cyst for *Malassezia*, and *Pseudophaeomoniella oleicola*. Of interest, the samples with higher probability of fungal taxa were the peritoneal fluids.

## Discussion

Herein, we report the results from a metagenomics community composition analysis in various tissue samples from a family of wild voles, one of which was affected with two extra-hepatic tapeworm cysts. The tapeworm identified, *H. taeniaeformis*, is widely abundant globally^27^. Like other tapeworms, this parasite has been documented in a variety of mammals, primarily infecting cats and other feline species^28^ (**Figure 4A**). *H. taeniaeformis* typically infects felines, having rodents serve as the primary intermediate host for the larval form, Strobilocercus^29,30^.

**Figure 4.**
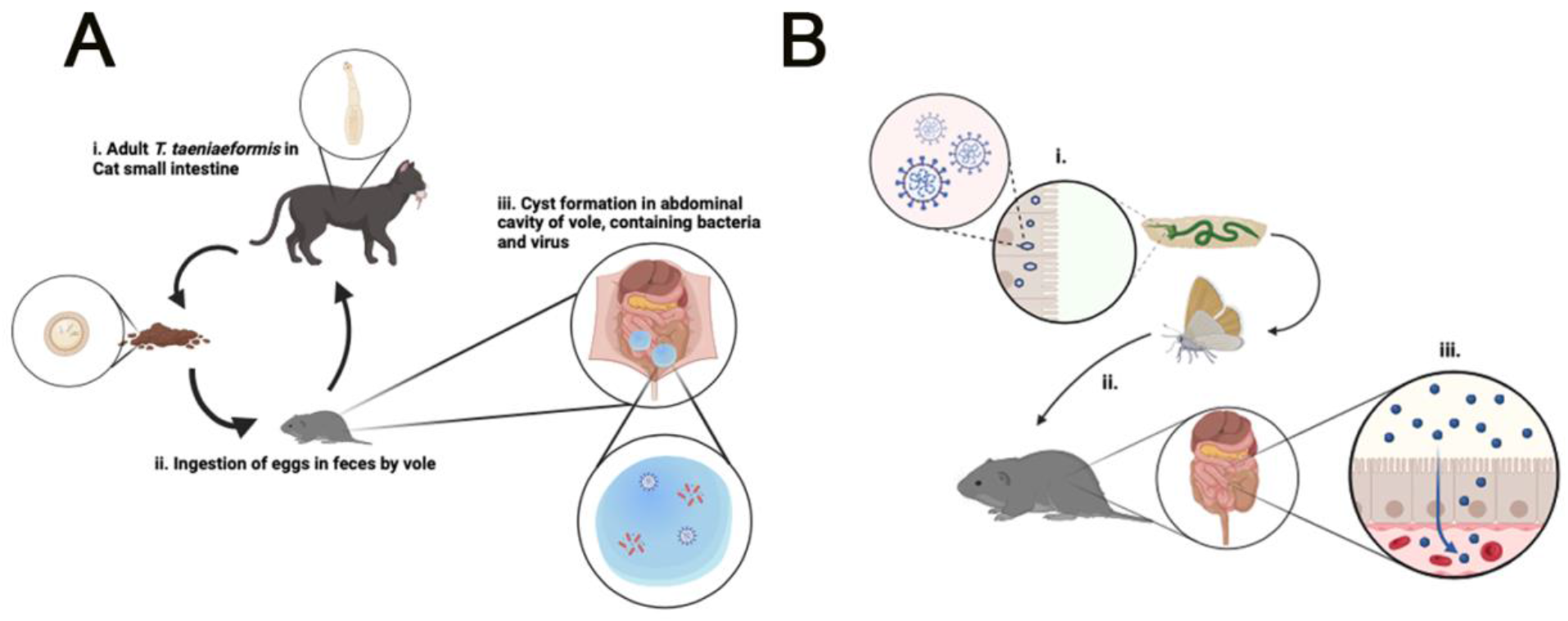
Overview of *Hydatigera taeniaeformis* lifecyle and metacestode larval cystic stages in rodents and our theory of how MyunGV-A viral DNA could reach and co-adapt with bacteria in tapeworm cysts. **A)** Left panel. *H. taeniaeformis* responsible for cyst formation in the peritoneal cavity in rodents may serve as a conducive environment for the retention and replication of MyunGV-A and bacteria. The cysts may offer protection against external factors and create a microenvironment suitable for these microorganisms, facilitating evasion of the immune system. **B)** Right panel. Presumptive theory for acquisition of the insect virus by voles is presumed to have occurred indirectly through dietary sources e.g., ingestion of insects or vegetation carrying MyunGV-A contaminants. Steps in proposed cycle: i., MyunGV-A replicates in the midgut epithelial cells of armyworms; ii., vole consumes armyworm moth; iii., MyunGV-A is absorbed by via the brush border of the intestinal epithelial cells and into the bloodstream, disseminating to the cysts, or follows the *H. taeniaeformis* as it migrates through the gut wall to extra-intestinal tissues to start the cystic stage of the tapeworm.

From a microbiome perspective, it is well known that tapeworms affect the gut microbiome in humans and animals^15,31-33^, they produce secretory molecules which affect the gut microbiota^34^, and that the infestation promotes the production of immunoglobulins (IgG, IgG1, IgG2a, IgG2b, IgG3 and IgM) against gut commensals that correlate with increase or decrease in the feces^15^. Despite this knowledge, little is known about the microbiome features of the cystic structures of tapeworm larva or other migratory parasites and the potential they may have to cause local infections. In our study, the identification of bacteria inside the parasitic peri-hepatic cysts, opens new possibilities for understanding the complicated interplay of viruses, parasites, and bacteria in the peritoneal cavity of mice.

From an ecological perspective, the unanticipated discovery of Mythimna unipuncta granulovirus A (MyunGV-A), commonly a lepidopteran-specific baculovirus, in addition to a community of bacteria such as *Lactobacillus acidophilus, Enterococcus faecium, Bacteroidales*, and *Klebsiella variicola*, inside *H. taeniaeformis* cysts in wild voles challenges conventional theories of host-virus specificity and microbial dynamics in ecosystems. Although we did not verify the presence of viruses with cultivation methods or electron microscopy visualization, the presence of MyunGV-A is intriguing because Baculoviruses, such as MyunGV-A, use receptors on gut cells of their insect hosts to enable the virus entry, reproduction, and dissemination within the insect cells^35,36^. The gastrointestinal system of voles, which is normally a reservoir for a diverse range of commensal bacteria^37,38^ could serve as a suitable environment for MyunGV-A survival and likely replication which we did not visualize^39^ or quantify^40^. It is unclear if the virus interacts with the strobilocercus microbiota, potentially aiding the survival of the community within the parasite in the mammalian host. If it is not adaptation with replication, another possibility to observe the infection of the mouse could be through dietary acquisition, in which voles consume MyunGV-A-carrying insects and temporarily allow the virus to replicate, until the host clears the viral infection. Although the virus does not reproduce within the vole digestive tract, it is possible that it could survive in the gastrointestinal environment, if the virus symbiosis takes place with bacteria, which has not been reported in MyunGV-A viral laboratory strains. Additionally, the virus and bacteria could enter the systemic circulation via breaks in the mucosal surfaces of the digestive tract of the voles, resulting in broad translocation and colonization throughout the peritoneal cavity. The specificity of these viruses is highlighted by their successful usage as biopesticides^41,42^, which is due to their inability to cross species and infect non-target mammals. Therefore, the existence of MyunGV-A within the rodent tapeworm larval cysts needs further examination before considering this presumed co-adaptation in migratory parasites as a strategy for non-traditional viral transmission and survival mechanisms outside the insect gut.

The findings also complement the results from a recent study where parasitic tapeworms of domestic animals in China showed a highly variable virome, but did not identify MyunGV-A in adult parasites^43^ which indicated several possibilities and theories for future testing (**Figure 4B**). Those theories include that adult tapeworm of domestic animals do not carry MyunGV-A or that there are virome differences in China vs USA with China not having MyunGV-A, or that MyunGV-A is likely associated with the gut microbiome of wild voles locally in Ohio being also independent of the viral load of adult tapeworms.

In conclusion, our findings revealed a simplified microbial community within the parasitic cysts of wild rodents, which contained gut commensals (*P. distasonis*) and an unexpected insect virus, which thrive reproducibly and independently in spatially-restricted niches, away from the gut, devoid of host-nutrient availability from ingested food or gut ingesta. Metacestodes represent a unique model of bacterial community evasion of the immune system and survival in a host-nutrient-deprived, parasite-acquired environment. This study provides a new perspective on the understanding of bacterial communities in migratory tapeworms and in dysbiosis associated with chronic intestinal diseases where spatially-restricted cavitating micro niches develop and could perpetuate inflammation through symbiotic mechanisms. Such a scenario is in Crohn’s disease, where recently we discovered that *P. distasonis* predisposes susceptible hosts to inflammation driven by succinate (*P. distasonis* metabolite)^7^, which increases cytotoxicity of immune cells to co-habiting *Escherichia coli*. Our study could help identify revised strategies for the treatment of clinical cysticercosis and the potential benefits of adding antibiotics.

## Acknowledgements

This project was conducted with discretionary funds to support the NIH study grant R21 DK118373 to A.R.-P., entitled “Identification of pathogenic bacteria in Crohn’s disease.” We thank Afnan Khan and Brandon Grubb for their scientific contributions.

## References

1. Webster NS. Cooperation, communication, and co-evolution: grand challenges in microbial symbiosis research. Front Microbiol. 2014;5:164. doi:10.3389/fmicb.2014.00164

2. Raina JB, Eme L, Pollock FJ, Spang A, Archibald JM, Williams TA. Symbiosis in the microbial world: from ecology to genome evolution. Biol Open. Feb 22 2018;7(2)doi:10.1242/bio.032524

3. Yeoman CJ, Chia N, Yildirim S, et al. Towards an Evolutionary Model of Animal-Associated Microbiomes. Entropy. 2011;13(3):570–594.

4. McFall-Ngai MJ. Giving microbes their due--animal life in a microbially dominant world. J Exp Biol. Jun 2015;218(Pt 12):1968–73. doi:10.1242/jeb.115121

5. Rodriguez-Palacios A, Kodani T, Kaydo L, et al. Stereomicroscopic 3D-pattern profiling of murine and human intestinal inflammation reveals unique structural phenotypes. Nat Commun. 2015;6:7577. doi:10.1038/ncomms8577

6. Yang F, Kumar A, Davenport KW, et al. Complete Genome Sequence of a Parabacteroides distasonis Strain (CavFT hAR46) Isolated from a Gut Wall-Cavitating Microlesion in a Patient with Severe Crohn’s Disease. Microbiol Resour Announc. Sep 2019;8(36)doi:10.1128/MRA.00585-19

7. Singh V, West G, Fiocchi C, et al. Clonal Parabacteroides from Gut Microfistulous Tracts as Transmissible Cytotoxic Succinate-Commensal Model of Crohn’s Disease Complications. bioRxiv. Jan 10 2024;doi:10.1101/2024.01.09.574896

8. Bank NC, Vaidhvi S, Grubb B, et al. The basis of antigenic operon fragmentation in Bacteroidota and commensalism. bioRxiv 20230602543472; doi: 10.1101/20230602543472. 2023;

9. Singh V, Rodriguez-Palacios A. Genomes of Bacteroides ovatus, B. cellulosilyticus, B. uniformis, Phocaeicola vulgatus, and P. dorei isolated from Gut Cavernous Fistulous Tract Micropathologies in Crohn’s disease. MRA. 2024;

10. Caira JN, Jensen K. Electron microscopy reveals novel external specialized organs housing bacteria in eagle ray tapeworms. PLoS One. 2021;16(1):e0244586. doi:10.1371/journal.pone.0244586

11. Bonelli P, Serra E, Dei Giudici S, et al. Molecular phylogenetic analysis of Echinococcus granulosus sensu lato infecting sheep in Italy. Acta Trop. Apr 2024;252:107151. doi:10.1016/j.actatropica.2024.107151

12. Romig T, Wassermann M. Echinococcus species in wildlife. Int J Parasitol Parasites Wildl. Apr 2024;23:100913. doi:10.1016/j.ijppaw.2024.100913

13. Celik F, Selcuk MA, Kilinc SG, et al. Molecular discrimination of G1 and G3 genotypes of Echinococcus granulosus sensu stricto obtained from human, cattle, and sheep using the mitochondrial NADH dehydrogenase subunit 5 marker. Acta Trop. Apr 2024;252:107124. doi:10.1016/j.actatropica.2024.107124

14. Casulli A, Pane S, Randi F, et al. Primary cerebral cystic echinococcosis in a child from Roman countryside: Source attribution and scoping review of cases from the literature. PLoS Negl Trop Dis. Sep 2023;17(9):e0011612. doi:10.1371/journal.pntd.0011612

15. Bao J, Zheng H, Wang Y, et al. Echinococcus granulosus Infection Results in an Increase in Eisenbergiella and Parabacteroides Genera in the Gut of Mice. Front Microbiol. 2018;9:2890. doi:10.3389/fmicb.2018.02890

16. Browne HP, Forster SC, Anonye BO, et al. Culturing of ‘unculturable’ human microbiota reveals novel taxa and extensive sporulation. Nature. 05 2016;533(7604):543–546. doi:10.1038/nature17645

17. Szabó BG, Kiss R, Makra N, et al. Composition and changes of blood microbiota in adult patients with community-acquired sepsis: A. Front Cell Infect Microbiol. 2022;12:1067476. doi:10.3389/fcimb.2022.1067476

18. Thoendel M, Jeraldo P, Greenwood-Quaintance KE, et al. Comparison of Three Commercial Tools for Metagenomic Shotgun Sequencing Analysis. J Clin Microbiol. Feb 24 2020;58(3)doi:10.1128/JCM.00981-19

19. CosmosID. Metagenomics Cloud, app.cosmosid.com, CosmosID Inc., https://www.cosmosid.com 2024;

20. Yan Q, Wi YM, Thoendel MJ, et al. Evaluation of the CosmosID Bioinformatics Platform for Prosthetic Joint-Associated Sonicate Fluid Shotgun Metagenomic Data Analysis. J Clin Microbiol. Feb 2019;57(2) doi:10.1128/JCM.01182-18

21. Martin W. Linking causal concepts, study design, analysis and inference in support of one epidemiology for population health. Preventive Veterinary Medicine. 2008;86(3-4):270–288.

22. Rodriguez-Palacios A, Aladyshkina N, Ezeji JC, et al. ‘Cyclical Bias’ in Microbiome Research Revealed by A Portable Germ-Free Housing System Using Nested Isolation. Sci Rep. Feb 2018;8(1):3801. doi:10.1038/s41598-018-20742-1

23. Raffner Basson A, Gomez-Nguyen A, LaSalla A, et al. Replacing Animal Protein with Soy-Pea Protein in an “American Diet” Controls Murine Crohn Disease-Like Ileitis Regardless of Firmicutes: Bacteroidetes Ratio. J Nutr. 03 2021;151(3):579–590. doi:10.1093/jn/nxaa386

24. Basson AR, Gomez-Nguyen A, Menghini P, et al. Human Gut Microbiome Transplantation in Ileitis Prone Mice: A Tool for the Functional Characterization of the Microbiota in Inflammatory Bowel Disease Patients. Inflamm Bowel Dis. Feb 11 2020;26(3):347–359. doi:10.1093/ibd/izz242

25. Harrison RL, Mowery JD, Bauchan GR, Theilmann DA, Erlandson MA. The complete genome sequence of a second alphabaculovirus from the true armyworm, Mythimna unipuncta: implications for baculovirus phylogeny and host specificity. Virus Genes. Feb 2019;55(1):104–116. doi:10.1007/s11262-018-1615-7

26. García M, Ortego F, Hernández-Crespo P, Farinós GP, Castañera P. Inheritance, fitness costs, incomplete resistance and feeding preferences in a laboratory-selected MON810-resistant strain of the true armyworm Mythimna unipuncta. Pest Manag Sci. Dec 2015;71(12):1631–9. doi:10.1002/ps.3971

27. Gomez-Puerta LA, Vargas-Calla A, Garcia-Leandro M, et al. Identification of wild rodents as intermediate hosts for Hydatigera taeniaeformis in Peru. Parasitol Res. Aug 2023;122(8):1915–1921. doi:10.1007/s00436-023-07892-6

28. Jia W, Yan H, Lou Z, et al. Mitochondrial genes and genomes support a cryptic species of tapeworm within Taenia taeniaeformis. Acta Trop. Sep 2012;123(3):154–63. doi:10.1016/j.actatropica.2012.04.006

29. Cook RW, Trapp AL, Williams JF. Pathology of Taenia taeniaeformis infection in the rat: hepatic, lymph node and thymic changes. J Comp Pathol. Apr 1981;91(2):219–26. doi:10.1016/0021-9975(81)90026-8

30. Mahesh Kumar J, Reddy PL, Aparna V, et al. Strobilocercus fasciolaris infection with hepatic sarcoma and gastroenteropathy in a Wistar colony. Vet Parasitol. Nov 05 2006;141(3-4):362–7. doi:10.1016/j.vetpar.2006.05.029

31. Zhu M, Wang C, Yang S, et al. Alterations in Gut Microbiota Profiles of Mice Infected with Echinococcus granulosus sensu lato Microbiota Profiles of Mice Infected with E. granulosus s.l. Acta Parasitol. Dec 2022;67(4):1594–1602. doi:10.1007/s11686-022-00613-6

32. Cao D, Pang M, Wu D, et al. Alterations in the Gut Microbiota of Tibetan Patients With Echinococcosis. Front Microbiol. 2022;13:860909. doi:10.3389/fmicb.2022.860909

33. Liu Z, Yin B. Alterations in the Gut Microbial Composition and Diversity of Tibetan Sheep Infected With. Front Vet Sci. 2021;8:778789. doi:10.3389/fvets.2021.778789

34. Wu J, Zhu Y, Zhou L, et al. Parasite-Derived Excretory-Secretory Products Alleviate Gut Microbiota Dysbiosis and Improve Cognitive Impairment Induced by a High-Fat Diet. Front Immunol. 2021;12:710513. doi:10.3389/fimmu.2021.710513

35. Clem RJ, Passarelli AL. Baculoviruses: sophisticated pathogens of insects. PLoS Pathog. 2013;9(11):e1003729. doi:10.1371/journal.ppat.1003729

36. Rohrmann GF. Baculovirus Molecular Biology. 2013.

37. Weldon L, Abolins S, Lenzi L, Bourne C, Riley EM, Viney M. The Gut Microbiota of Wild Mice. PLoS One. 2015;10(8):e0134643. doi:10.1371/journal.pone.0134643

38. Viney M. The gut microbiota of wild rodents: Challenges and opportunities. Lab Anim. Jun 2019;53(3):252–258. doi:10.1177/0023677218787538

39. Li Y, Liu X, Tang P, Zhang H, Qin Q, Zhang Z. Genome sequence and organization of the Mythimna (formerly Pseudaletia) unipuncta granulovirus Hawaiian strain. Sci Rep. Jan 11 2021;11(1):414. doi:10.1038/s41598-020-80117-3

40. Mukawa S, Goto C. In vivo characterization of two granuloviruses in larvae of Mythimna separata (Lepidoptera: Noctuidae). J Gen Virol. Apr 2008;89(Pt 4):915–921. doi:10.1099/vir.0.83365-0

41. Lacey LA, Grzywacz D, Shapiro-Ilan DI, Frutos R, Brownbridge M, Goettel MS. Insect pathogens as biological control agents: Back to the future. J Invertebr Pathol. Nov 2015;132:1–41. doi:10.1016/j.jip.2015.07.009

42. van Oers MM, Pijlman GP, Vlak JM. Thirty years of baculovirus-insect cell protein expression: from dark horse to mainstream technology. J Gen Virol. Jan 2015;96(Pt 1):6–23. doi:10.1099/vir.0.067108-0

43. Zhang P, Zhang Y, Cao L, et al. A Diverse Virome Is Identified in Parasitic Flatworms of Domestic Animals in Xinjiang, China. Microbiol Spectr. Jun 15 2023;11(3):e0070223. doi:10.1128/spectrum.00702-23

